# Reported CCR5-∆32 deviation from Hardy-Weinberg equilibrium is explained by poor genotyping of rs62625034

**DOI:** 10.1101/791517

**Authors:** Yosuke Tanigawa, Manuel A. Rivas

**Affiliations:** Department of Biomedical Data Science, Stanford University, Stanford, CA, 94305

## Abstract

In the fall of 2018, news broke about a researcher from China who had used CRISPR gene editing to cause human babies to have a deletion in the CCR5 chemokine receptor, making them resistant to HIV infection. One of the numerous ethical concerns about this study is that the deletion may have other effects. Subsequently, Nature Medicine published a Brief Communications from Wei and Nielsen concluding that homozygotes for the CCR5-∆32 deletion have a survival probability to age 76 of 83.5% compared to 86.5% and 86.4% for the heterozygotes and the other homozygote, respectively, and that observed departures from Hardy Weinberg proportions also support selection operating on this allele^1^. In the study, Wei and Nielsen used a proxy variant, rs62625034 in their analysis. Here, we report that the reported CCR5-∆32 deviation from Hardy-Weinberg equilibrium (HWE) inferred by Wei and Nielsen can be explained by poor genotyping of rs62625034, the variant used for their analysis.

## Main

In medical genetics studies, data quality assessment and control steps are typically carried out to reduce potential biases that may be introduced in an association study^2^. It is an integral part of genetic analysis and scrutiny is always required of novel associations, which includes scrupulous assessment of intensity cluster plots of variants found to be associated and whether potential batch effects may be responsible for the putative signals. Errors in genotyping calling have the potential to introduce systematic biases, leading to an increase in the number of false-positive throughout the analytical workflow including estimates of allele frequency, estimates of deviation from HWE, and association statistics. Most genome-wide association studies choose to exclude markers that show extensive deviation from Hardy-Weinberg equilibrium (HWE) because this can be indicative of a genotyping or genotype calling error as we recently did in our association analysis of UK Biobank array data where we excluded over 50,000 variants from our analysis^3^. However, deviations from HWE may also indicate selection and it would be unwise to remove these loci from further genetic analysis. However, when the HWE thresholds are increased it is typical for researchers who find potential signals in their data to carefully examine all genotype cluster plots for SNPs showing some evidence of deviation from HWE manually.

Here, we systematically examine the genotyping performance of rs62625034 in white British individuals from UK Biobank (n = 409,634). The variant rs62625034 was genotyped using both the UK Biobank Axiom array (n = 95 batches, 2 failed genotyping) and UK BiLEVE^4^ (n = 11). Indeed, as in Wei and Neilsen, we also estimate deviation from HWE for rs62625034. Given the strong deviation from HWE and the high missingness rate (3.61%) estimated from genotypes made available by UK Biobank we proceeded with manually inspecting the cluster plots for rs62625034. We used ScatterShot from McCarthy group at Oxford University, which provides statistics and visualization of genotype clusters aggregated across all batches and separated by batch and genotyping array (Figure 1A). We found that the cluster plot provides poor support for a clear separation between the three genotyping classes. As a result, we next asked whether other proxy variants for rs333, the CCR5-∆32 deletion, exhibited similar poor genotyping performance. Using LDproxy from NCI and selecting GBR and CEU population from 1000 Genomes project we found that rs113010081 is a proxy variant (r=0.964) and genotyped in UK Biobank^5^. We found that the variant has a low missingness rate 0.08%, no deviation from Hardy-Weinberg equilibrium (p-value = 0.369, Figure 1E), and three clearly separated genotype clusters indicative of a properly genotyped variant (Figure 1B). Given these observations, we asked whether the deviation from HWE for rs62625034 could be explained by batches with poor genotyping performance. We aggregated genotyping rate, heterozygosity, batch identifier, array type, and per batch HWE p-value calculation for the white British individuals in UK Biobank (**Supplementary Table S1**). We found a significant association (p = 0.001) between batch genotyping rate and HWE p-value deviation with batches with higher missingness rate (lower genotyping rate) corresponding to higher −log10(HWE p-values) (Figure 1C). Separating the analysis between UK Biobank Axiom array and UK BiLEVE array clearly show that the deviation from HWE observed for rs62625034 is accounted for by poor genotyping in UK Biobank Axiom array (KS test p = 0.00059 for −log10(HWE p-value) comparison between batches with high [>= 98%] and low [< 98%] genotyping rates). Similarly, 0 of the 11 UK BiLEVE genotyping batches had deviation from HWE (all p > 0.01).

**Figure 1.**
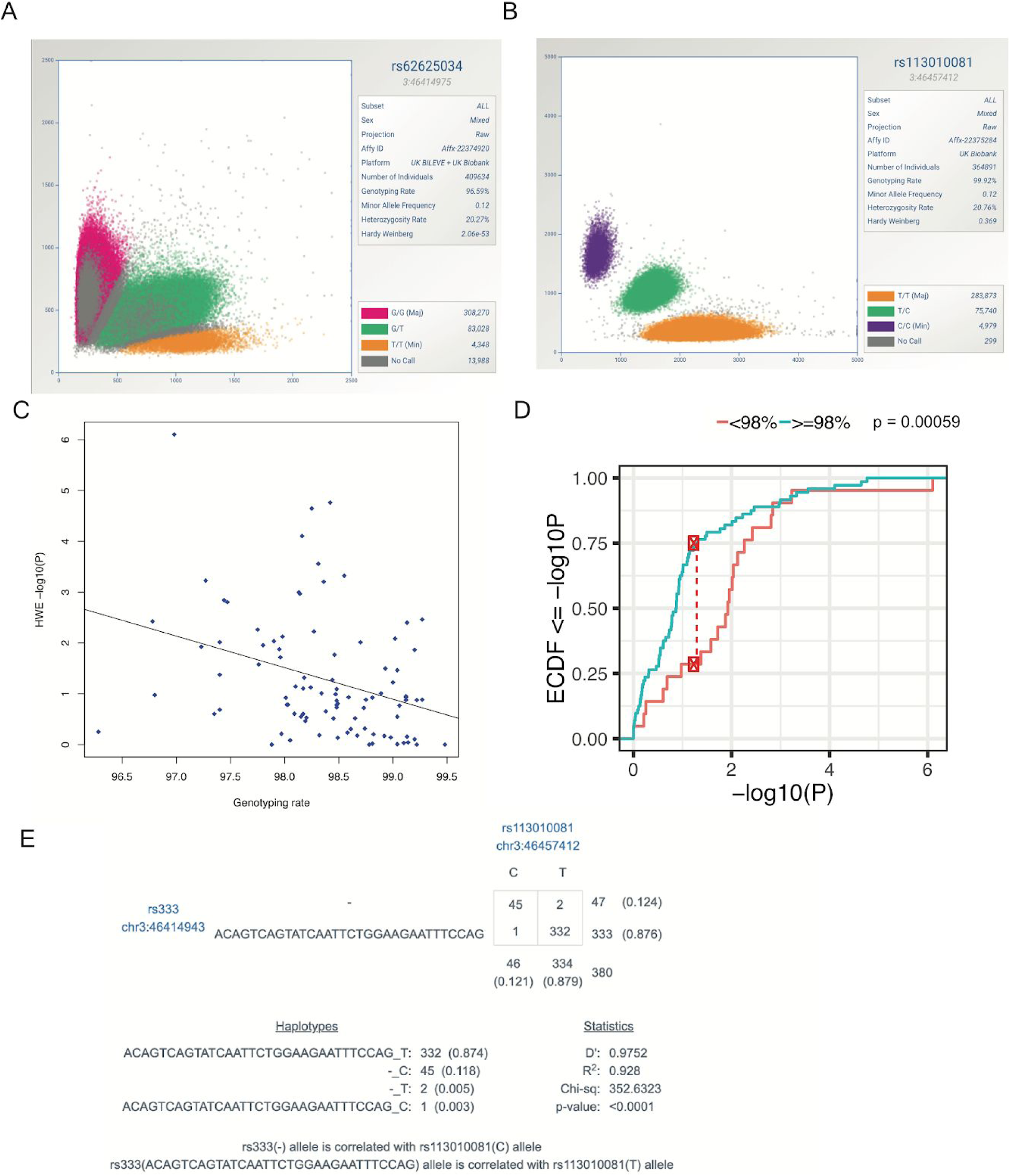
Poor genotyping for rs62625034 accounts for departure from HWE in UK Biobank. **A)** Cluster plot of three genotype classes for rs62625034 (the studied variant in Wei and Nielsen), magenta corresponds to G/G (homozygous reference allele genotype), green corresponds to G/T (heterozygous genotype), and T/T (homozygous alternate allele genotype) (source ScatterShot for White British individuals in UK Biobank^7^). Genotyping rate equal to 96.59% and HWE p-value = 2.06e-53. **B)** Cluster plot of three genotype classes for rs113010081 (a proxy variant for rs333, the CCR5-delta 32 variant). Genotyping rate of 99.92% and HWE p-value = 0.369 (HWE p-value = 0.16 for white-British unrelated individuals in UK Biobank as described in Tanigawa et al.^3^). **C)** Scatterplot of the genotyping rate (x-axis) versus Hardy-Weinberg Equilibrium deviation p-value (y-axis) for rs62625034. Each point represents a batch from the 95 batches in Axiom Biobank array (batch number 66 and 47 were excluded as genotype calls were generated). Best fit line is shown (p = .001). **D**) The cumulative distribution function of −log10(HWE p-value) for batches with genotyping rate above 98% and batches with genotyping rate below 98%. **E)** Linkage disequilibrium statistics for CEU/GBR European individuals in the 1000 Genomes project between rs333 (delta 32 variant) and rs113010081 (r = 0.964).

Together, these data show that careful analysis should be warranted when dealing with variants with patterns consistent with poor genotyping performance. The conclusions drawn in Wei and Nielsen have an alternative explanation to selection. Analogous to prior publications in the medical genetics community, we discovered that technical errors and inadequate quality control protocol introduced false-positive findings in Wei and Nielsen and that the deviation from HWE can be explained by poor genotyping performance^6^.

## Supplementary Information

**Supplementary Table S1**. rs62625034 genotyping rate, HWE deviation, and heterozygosity for each genotyping batch in UK Biobank.

